# Turnover of the extracellular polymeric matrix in an EBPR microbial community

**DOI:** 10.1101/2022.08.11.503576

**Authors:** Sergio Tomás-Martínez, Erwin J. Zwolsman, Franck Merlier, Martin Pabst, Yuemei Lin, Mark C.M. van Loosdrecht, David G. Weissbrodt

## Abstract

Polyphosphate accumulating organisms (PAOs) are responsible for enhanced biological phosphate removal (EBPR) from wastewater, where they grow embedded in a matrix of extracellular polymeric substances (EPS). EPS comprise a mixture of biopolymers like polysaccharides or (glyco)proteins. Despite previous studies, little is known about the dynamics of EPS in mixed cultures, and their production by PAOs and potential consumption by flanking microbes. EPS are biodegradable and have been suggested to be a substrate for other organisms in the community. Studying EPS turnover can help elucidate their biosynthesis and biodegradation cycles. We analyzed the turnover of proteins and polysaccharides in the EPS of an enrichment culture of PAOs relative to the turnover of internal proteins. An anaerobic-aerobic sequencing batch reactor (SBR) simulating EBPR conditions was operated to enrich for PAOs. After achieving a stable culture, carbon source was switched to uniformly ^13^C-labelled acetate. Samples were collected at the end of each aerobic phase. EPS were extracted by alkaline treatment. ^13^C enrichment in proteins and sugars (after hydrolysis of polysaccharides) in the extracted EPS were measured by mass spectrometry. The average turnover rate of sugars and proteins (0.167 and 0.192 d^-1^ respectively) was higher than the expected value based on the solid removal rate (0.132 d^-1^), and no significant difference was observed between intracellular and secreted proteins. This indicates that EPS from the PAO enriched community is not selectively degraded by flanking populations under stable EBPR process conditions. Instead, we observed general decay of biomass, which corresponds to a value of 0.048 d^-1^.

**Graphical abstract:** 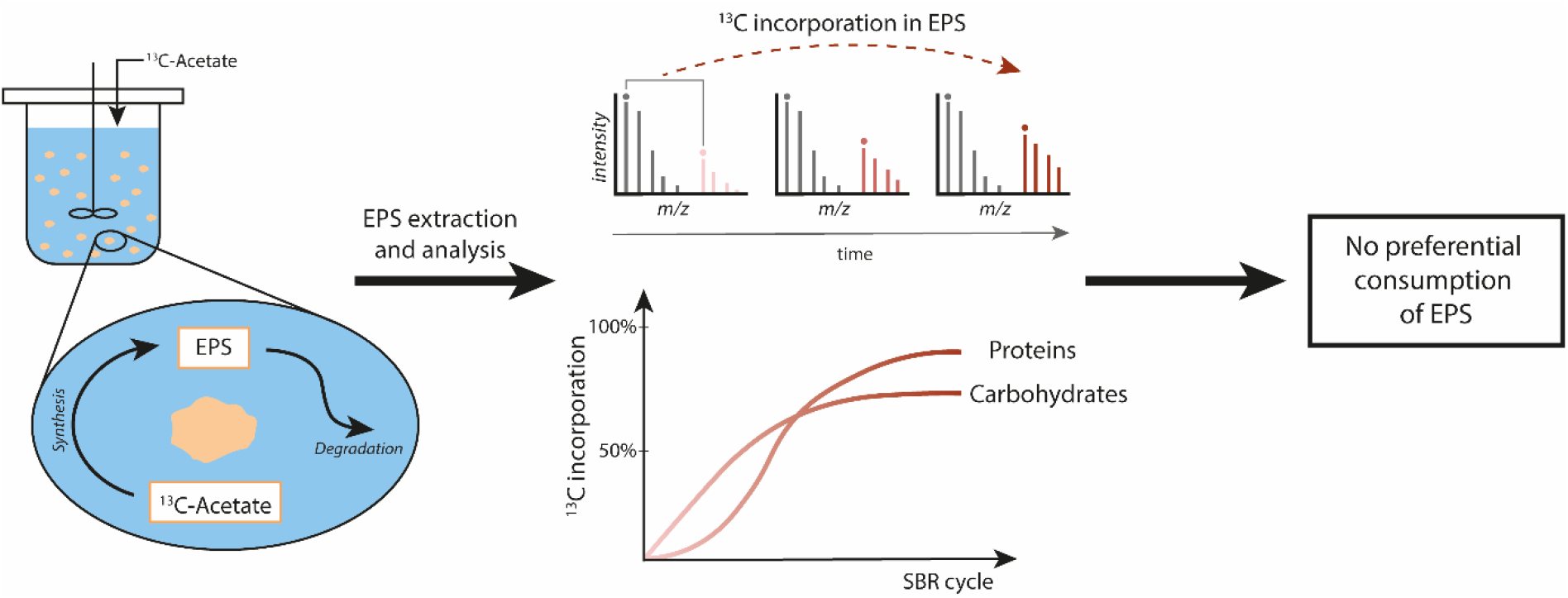

**Key points:**

- Proteins showed a higher turnover rate than carbohydrates.
- Turnover of EPS was similar to the turnover of intracellular proteins.
- EPS is not preferentially consumed by flanking populations.

## 1. Introduction

Enhanced biological phosphorus removal (EBPR) is a widely utilized treatment to eliminate inorganic phosphorus and organic matter from wastewater. Polyphosphate-accumulating organisms (PAOs) have the capacity to anaerobically take up volatile fatty acids (*e.g*., acetate) and to store them intracellularly as polyhydroxyalkanoate (PHA). When an electron donor is available, PAOs consume the stored PHA and take up inorganic phosphate, storing it as intracellular polyphosphate (Smolders et al. 1995). PAOs are a very important group of organisms in wastewater treatment plants performing EBPR, including the aerobic granular sludge process. *“Candidatus* Accumulibacter phosphatis” has been described as the dominant species responsible for phosphorus removal, although other PAOs have been identified, such as *Tetrasphaera* or *Dechloromonas* (Marques et al. 2017; Petriglieri et al. 2021). In wastewater treatment plants, these microorganisms grow as part of the microbial community in the form of bioaggregates (flocs, granules or biofilms) embedded in a matrix of extracellular polymeric substances (EPS) (Weissbrodt et al. 2013; Barr et al. 2016).

EPS are metabolically synthesized and released by the microorganisms forming scaffolds that provide mechanical stability in biofilms. EPS consist of a mixture of biopolymers, such as polysaccharides, proteins, nucleic acids or lipids, among others (Flemming and Wingender 2010). The EPS matrix provides certain benefits to the microbial community such as water retention, nutrient sorption or protection against viruses and predators (Seviour et al. 2019). Therefore, it is important to keep the extracellular structure stable.

EPS is regularly indicated as a potential nutrient source for the microorganisms present in the community (Flemming and Wingender 2010). A few studies have shown the biodegradability of EPS, *e.g*., Zhang and Bishop (2003) have demonstrated that extracted EPS can be consumed by their own producers when supplied as substrate. The same results have been obtained by Wang, Liu and Tay (2007) using extracted EPS from aerobic granules. Pannard et al. (2016) have shown the degradation of EPS under nitrogen and carbon limitation. Some microorganisms present in the EBPR systems, *e.g*., *Sphingobacterium* or *Chryseobacterium*, have been described to consume exopolysaccharides and other complex polymeric compounds (Matsuyama et al. 2008; McBride 2014). Therefore, granules could be seen as an ecosystem containing EPS producers and consumers (Weissbrodt et al. 2013). Despite that the biodegradability of EPS has been proven, the consumption has been only shown under substrate limited conditions and using extracted EPS (*ex situ*). The *in situ* consumption of EPS under stable culture conditions has not been shown, although often implicit or explicit assumed to occur based on the above described type of experiments. Therefore, the consumption of EPS by flanking populations (*i.e*., low abundant bacteria that do not grow on the primary substrate) needs to be evaluated to prove these assumptions. During normal growth conditions, the consumption of a certain compound or polymer needs to be balanced by the production of newly synthesized ones, to maintain the steady state. This renewal or replacement is referred as turnover, and the frequency at which it happens, as turnover rate (Doherty and Whitfield 2011).

In order to evaluate the turnover of biopolymers, newly synthesized polymers have to be differentiated from the already existing ones. Metabolic incorporation of a stable isotope (*e.g*., ^13^C or ^15^N) is a widely used technique for the study of the dynamics of different polymers. Although most of the research focuses in the isotope incorporation in proteins (Jehmlich et al. 2016), some studies have also focused on the incorporation into carbohydrates or lipids (Liang et al. 2017; Schlame et al. 2020). In some pure cultures, the use of labelled building blocks (*e.g*., a specific amino acid or monosaccharide) is preferred. However, this is not suitable for most organisms, especially for microbial communities, where these building blocks can be used for other purposes rather than anabolism. In these cases, the use of a labelled substrate is preferred (*e.g*., ^13^C-labelled carbon source) (Taubert et al. 2011). This metabolic labelling technique does not limit the study to a certain polymer (*e.g*., the use of labelled amino acids can only be used for the study of proteins), but allows to study all microbial products as the stable isotope is incorporated through the central metabolism (Lawson et al. 2021). Label incorporation can be analyzed by mass spectrometric techniques obtaining parameters such as the fraction of stable isotope incorporation (relative isotope abundance, RIA) and labelling ratio (LR). The LR describes the ratio of the labelled compound to the non-labeled compound and can be used for the estimation of the turnover rate of biomolecules (Jehmlich et al. 2016).

Several studies have applied stable isotope labelling for the analysis of EPS metabolism (Moerdijk-Poortvliet et al. 2018; Maqbool et al. 2020; Arshad et al. 2021). Specifically, Maqbool et al. (2020) employed ^13^C and ^15^ N labelled substrates to evaluate the turnover of the EPS of activated sludge. Although they have demonstrated that carbon content of the EPS is replenished faster than nitrogen, specific components such as proteins or polysaccharides were not analyzed. Additionally, Arshad et al. (2021) have shown no difference between the carbon and nitrogen assimilation to biomass or to EPS. Since the main components of the EPS contain carbon and nitrogen in their chemical structures, such as amino acids, amino-sugars or sialic acids, it is not possible to derive conclusions on the different specific components of the EPS.

The aim of the current research is to evaluate the turnover dynamics of the main components of the EPS of PAOs (*i.e*., polysaccharides and proteins) in comparison with the intracellular polymer turnover. We evaluated the specific label (*i.e*., ^13^C) incorporation patterns in the proteins and polysaccharides. A sequencing batch reactor (SBR) simulating EBPR conditions was operated to enrich a PAO dominated microbial community. Once stable conditions were achieved, the enrichment was fed with uniformly ^13^C-labelled acetate over one solid retention time (SRT). Samples were collected at the end of each SBR cycle and EPS were extracted and incorporation of ^13^C in proteins and sugar monomers (after the hydrolysis of the original polysaccharides) from the extracted EPS was measured by mass spectrometry. Sequence prediction tools were used to differentiate between intracellular and extracellular proteins.

## 2. Materials and methods

### 2.1. Sequencing batch reactor operation

The PAO enrichment was obtained in a 1.5 L (1 L working volume) sequencing batch reactor (SBR), following conditions similar to the one described by Guedes da Silva et al. (2020) with some adaptations. The reactor was inoculated using activated sludge from a municipal wastewater treatment plant (Harnaschpolder, The Netherlands). Each SBR cycle lasted 6 hours, consisting of 20 minutes of settling, 15 minutes of effluent removal, 5 minutes of N_2_ sparging, 5 minutes of feeding, 135 minutes of anaerobic phase and 180 minutes of aerobic phase. The hydraulic retention time (HRT) was 12 hours (removal of 500 mL of broth per cycle). The average solids retention time (SRT) was controlled to 7.55 ± 0.15 days by the removal of effluent at the end of the mixed aerobic phase. The pH was controlled at 7.4 ± 0.1 by dosing 0.2 M HCl or 0.2 M NaOH. The temperature was maintained at 20 ± 1 ^o^C.

The reactor was fed with two separate media: a concentrated COD medium (8.51 g/L NaAc·3H_2_O, 0.04 g/L yeast extract) and a concentrated mineral medium (1.53 g/L NH_4_Cl, 1.59 g/L MgSO_4_·7H_2_O, 0.40 g/L CaCl_2_·2H_2_O, 0.48 KCl, 0.04 g/L N-allylthiourea (ATU), 2.22 g/L NaH_2_PO_4_·H_2_O, 6 mL/L of trace element solution prepared as (Smolders et al. 1994)). In each cycle 50 mL of each media was added to the reactor, together with 400 mL of demineralized water. The final feed contained 400 mg COD/L of acetate.

#### 2.1.1. Monitoring of the SBR

Conductivity in the bulk liquid was used to follow phosphate release and uptake patterns and to verify the steady performance of the reactor. Extracellular concentrations of phosphate and ammonium were measured with a Thermo Fisher Gallery Discrete Analyzer (Thermo Fisher Scientific, Waltham, MA). Acetate was measured by high performance liquid chromatography (HPLC) with an Aminex HPX-87H column (Bio-Rad, Hercules, CA), coupled to an RI and UV detectors (Waters, Milford, MA), using 0.01 M phosphoric acid as eluent supplied at a flowrate of 0.6 mL/min.

### 2.2. Microbial community characterization

#### 2.2.1. Fluorescence in situ hybridization (FISH)

Samples were handled, fixed and stained as described by Winkler et al. (2011). All bacteria were targeted using a mixture of EUB338, EUB338-II and EUB338-III probes (Amann et al. 1990; Daims et al. 1999). “*Ca*. Accumulibacter” was visualized using a mixture of PAO462, PAO651 and PAO846 probes (PAOmix) (Crocetti et al. 2000). *Dechloromonas* was targeted using the probe Dech69 and Dech209 (Dechmix). Hybridized samples were examined with Axio Imager 2 fluorescence microscope (Zeiss, Oberkochen, Germany).

#### 2.2.2. 16S rRNA gene amplicon sequencing

DNA was extracted from the granules using the DNeasy UltraClean Microbial kit (Qiagen, Venlo, The Netherlands), using the manufacturer’s protocol. The extracted DNA was quantified using a Qubit 4 (Thermo Fisher Scientific, Waltham, MA). Samples were sent to Novogene Ltd. (Hong Kong, China) for amplicon sequencing of the V3-4 hypervariable region of the 16S rRNA gene (position 341-806) on a MiSeq desktop sequencing platform (Illumina, San Diego, CA) operated under paired-end mode. The raw sequencing reads were processed by Novogene Ltd. (Hong Kong, China) and quality filtered using the QIIME software (Caporaso et al. 2010). Chimeric sequences were removed using UCHIME (Edgar et al. 2011) and sequences with ≥97% identity were assigned to the same operational taxonomic units (OTUs) using UPARSE (Edgar 2013). Each OTU was taxonomically annotated using the Mothur software against the SSU rRNA database of the SILVA Database (Quast et al. 2013).

#### 2.2.3. Metagenome sequencing

The previous DNA samples were sent to Novogene Ltd. (Hong Kong, China) for shotgun metagenome sequencing. DNA sample was fragmented by sonication to a size of 350 bp and the fragmented DNA was used for libraries preparation using NEB Next Ultra DNA Library Prep Kit, following manufacturer’s recommendations. Libraries were sequenced on an Illumina NovaSeq platform (Illumina, San Diego, CA) as 2 x 150 bp paired-end reads. The raw sequencing reads were processed by Novogene Ldt. (Hong Kong, China) for quality-trimming, adapter removal and contaminant-filtering. The trimmed reads were assembled into scaffolds using MEGAHIT (Li et al. 2015). Scaffolds (>=500 bp) were used for open reading frame (ORF) prediction by MetaGeneMark (Zhu et al. 2010). KEGG, EggNog and CAZy databases were used for functional annotation. Additionally, BLASTp from the NCBI website (blast.ncbi.nlm.nih.gov/Blast.cgi) for functional and taxonomic annotation. SignalP5.0 was employed for prediction of secreted proteins (Almagro Armenteros et al. 2019).

#### 2.2.4. Proteomic composition

Proteomic data annotated with taxonomies from an unlabeled sample (prior to the addition of ^13^C-acetate) was used to estimate the composition of the microbial community. Taxonomic composition was estimated based on the combined peptide areas corresponding to each genus.

### 2.3. ^13^C labelling experiment

After a stable enrichment was achieved, the feeding media was replaced in order to perform the labelling experiment. Some changes in the media were made for the experiment: yeast extract was added to the mineral media and COD media was prepared using only uniformly labelled sodium U-^13^C-acetate (99 atom % ^13^C, Cortecnet, Les Ulis, France), to a concentration of 5.25 g/L. Online conductivity measurements were used to ensure the correct performance of the enrichment during the experiment. These conditions were maintained during 8 days (corresponding to 32 cycles). Biomass samples were manually collected at the end of each aerobic phase maintaining the SRT. Collected biomass samples were washed and stored at - 80 °C.

### 2.4. EPS extraction and general analysis

Biomass samples were freeze-dried prior to the EPS extraction. EPS were extracted in alkaline conditions at high temperature, using a method adapted from Felz et al. (2016). Around 150 mg of freeze-dried biomass were stirred in 10 mL of 0.1 M NaOH (1.5 % w/v) at 80 °C for 30 min. Extraction mixtures were centrifuged at 4000xg at 4 °C for 20 min. Supernatants were collected and dialyzed overnight in dialysis tubing with a molecular cut-off of 3.5 kDa, frozen at −80 °C and freeze-dried. The freeze-dried extracted EPS samples were stored for further analysis.

The organic and ash fraction of the extracted EPS samples were determined by combusting them at 550 °C (APHA 1998). Protein content was estimated using the bicinchoninic acid (BCA) assay (Smith et al. 1985) with bovine serum albumin (BSA) as standard. Saccharides content was determined using the phenol-sulfuric acid assay (Dubois et al. 1956) with glucose as standard. Both methods were used as described by Felz et al. (2019).

### 2.5. Analysis of ^13^C incorporation into polysaccharides

Extracted EPS samples were hydrolyzed in 1 M HCl with a sample concentration of 10 g/L. Hydrolysis was performed at 105 °C during 8 hours with occasional mixing. After hydrolysis samples were neutralized with 1 M NaOH. Samples were centrifuged at 13300 rpm for 5 min and the supernatant was filtered through a 0.22 μm PVDF filter. Quantification of the enrichment of sugars was performed by liquid chromatography-high resolution mass spectrometry (LC-HRMS) on an HPLC Agilent 1290 with DAD connected to Agilent Q-TOF 6538. HPLC was carried out on an Agilent Poroshell 120 HILIC-Z (100 x 2.1 mm ID, 2.7 μm) column connected to an Agilent Infinity 1290 HPLC kept at 40°C. The solvent system was A: 25 mM of ammonium formate in H2O adjusted at pH 11 with ammonium hydroxide and B: Acetonitrile. The gradient program began with 97 % B, then ramped to 89 % B at 12 min, for decreased to 80 % B in 8 min, and to 30 % in 5 minutes returned to the initial conditions and kept constant for 3 min. The flow-rate was 0.500 mL/min and injection volume is 5 μL. All compounds response were measured in ESI- and calibrated externally. The ESI Gas Temp is 200°C, Vcap −3000V, Drying Gaz was set at 10 L/min and Nebuliser at 30 psig. Fragmentor was set at 100 V. HRMS spectrum was registered at 2 Hz in the mass range of 60 to 1100 m/z with internal calibration. MassHunter software was used data processing. The isotopic ions abundances were extracted from the extracted ions chromatogram calculate from the monoisotopic ions [M-H]^-^ or [M-COO]^-^ peak. The average of both values was used for the analysis.

### 2.6. Analysis of ^13^C incorporation into proteins

Briefly, biomass material was disrupted using beads beating in B-PER reagent (Thermo Fisher Scientific, Waltham, MA)/TEAB buffer (50 mM TEAB, 1 % (w/w) NaDOC, adjusted to pH 8.0) buffer. The cell debris was further pelleted and the proteins were precipitated using ice cold acetone. The protein pellet was redissolved in 200 mM ammonium bicarbonate, reduced using DTT and alkylated using iodoacetamide and digested using sequencing grade Trypsin (Promega, Madison, WI). Aliquots of *ca*. 100 ng protein digest were further analyzed by a one dimensional shotgun proteomics approach using an ESAY nano LC 1200 coupled to a QE plus Orbitrap mass spectrometer (Thermo Fisher Scientific, Waltham, MA) using methods described recently (Lawson et al. 2021; Kleikamp et al. 2021). Raw data were analyzed using PEAKS Studio (Bioinformatics Solutions Inc., Waterloo, Canada) for determining the community composition and MetaProSIP (OpenMS, University of Tubingen, Germany) (Sachsenberg et al. 2015) integrated into the KNIME 4.0.1 analytics platform as described recently (Lawson et al. 2021), was used to determine ^13^C stable isotope incorporation into the proteome. The most abundant ^13^C enriched isotope peak cluster (compared to the native peptide peak) was used to represent the labelling ratio of each time point.. All peptide spectra were matched against a protein database generated from predicted open reading frames from the total metagenomic assembly.

### 2.7. Calculation of turnover rates

The evolution of the labelling ratio (LR) of the different compounds during the cultivation time was evaluated in order to estimate the different turnover rates. The label incorporation in sugars followed an exponential cumulative dynamic, described by Eq. 1. Non-linear regression was used to calculate the different parameters: “a”, which represents the asymptotic value of the curve, and “k” (d^-1^), which is the turnover rate.

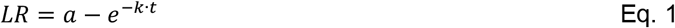

Proteins followed a sigmoidal dynamic, which can be described by the Gompertz function (Taubert et al. 2011). However, in order to compare the behavior of proteins to sugars, LR of proteins were also fitted using Eq. 1. Interestingly, the turnover rate calculated using Eq. 1 was similar to the one using the Gompertz function.

## 3. Results

### 3.1. PAO enrichment culture

In this study we analyzed the protein and polysaccharide turnover of a PAO enrichment, that was continuously cultivated in a sequencing batch reactor fed with acetate. A stable enrichment was confirmed by online pH and conductivity measurements as well as off-line measurements of acetate, phosphate and ammonium. The performance of a typical SBR cycle is shown in Fig. 1. Acetate was completely consumed anaerobically within the first 32 minutes, meaning that for most part of the cycle the only potential external carbon sources are EPS or other side products. Acetate consumption was linked to a net phosphate release of 134.2 mg PO_4_-P/L was achieved. This anaerobic release corresponds to 0.76 P-mol/C-mol of anaerobic phosphate release per carbon uptake. This is similar to previous highly enriched PAO communities (Smolders et al. 1994; Oehmen et al. 2005). During the aerobic phase, a phosphate removal of 207.9 mg PO_4_-P/L was achieved, resulting in a final phosphate concentration of 2.2 mg PO_4_-P/L. The decreasing ammonium concentration (7.2 mg NH_4_-N/L) was only associated to ammonium uptake for biomass growth, as nitrification was inhibited by the addition of ATU.

**Fig. 1.**
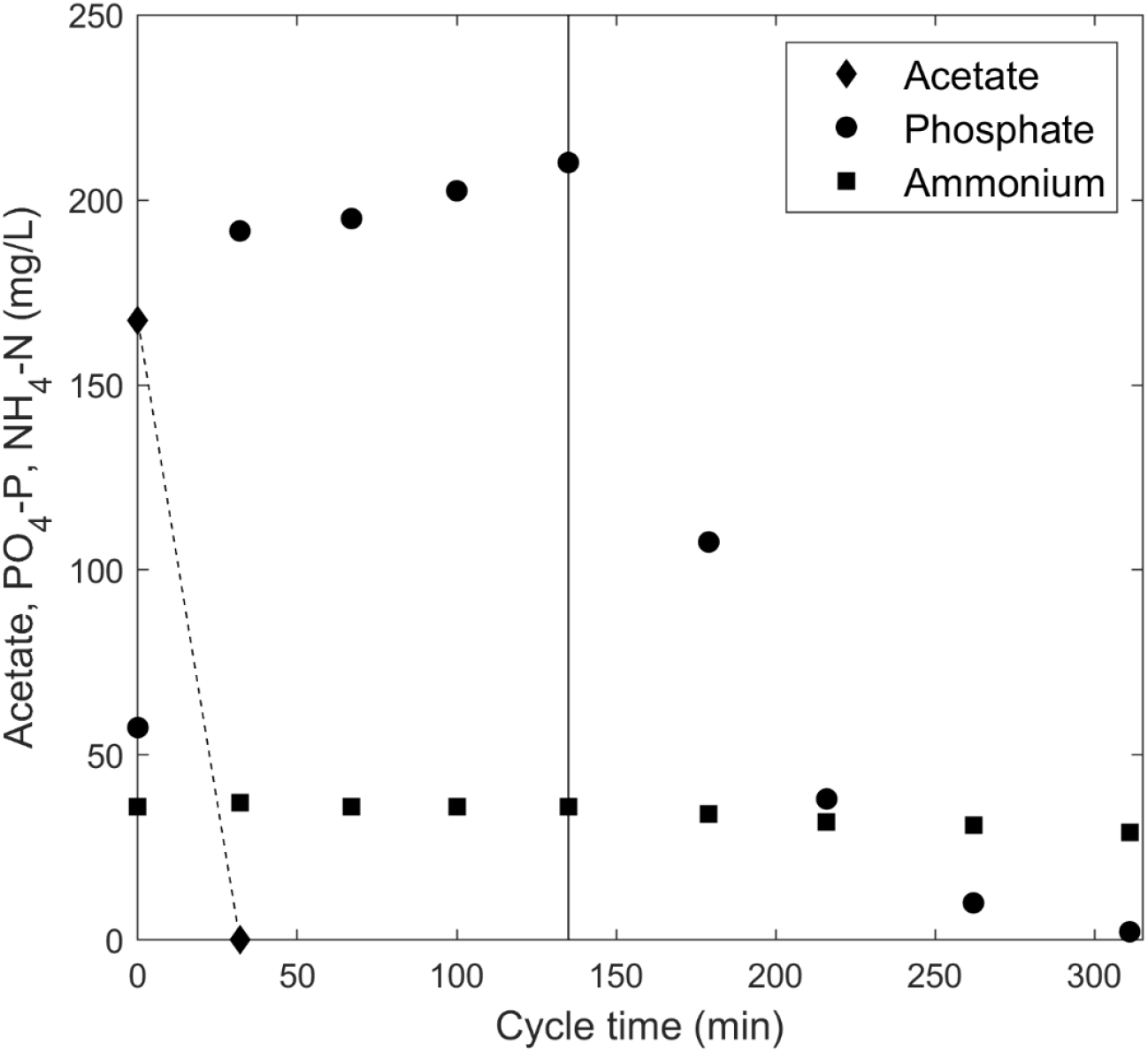
Concentration of acetate, phosphate and ammonium during a single SBR cycle after feeding. Black line represents the transition from anaerobic to aerobic phase

After a stable enrichment was obtained, the bacterial community composition was analyzed by means of FISH, 16S rRNA gene amplicon sequencing, and metaproteomic analysis (Fig. 2). Sequencing results showed a predominance of *Dechloromonas* (39.6 %) followed by the model PAO “*Ca*. Accumulibacter” (26.3 %). However, protein counts showed a higher predominance of *“Ca*. Accumulibacter” (60.7 %) compared to *Dechloromonas* (32.6 %). The microbial community compositions observed with these methods is further detailed in Fig. S1. Although glycogen accumulating organisms (GAOs) are typically seen in EBPR systems, they were not present in this microbial community. FISH results showed a large predominance of “*Ca*. Accumulibacter” relative to *Dechloromonas*. FISH was not exactly quantified but the ratio of “Ca. Accumulibacter” to *Dechloromonas* was roughly 10:1, while other eubacteria had a very low fraction in the community. Both populations of “*Ca*. Accumulibacter” and *Dechloromonas* have been described as PAO (Goel et al. 2012). Together with the reactor performance described in Fig. 1, the molecular data collectively demonstrate that a high PAO enrichment was achieved.

**Fig. 2.**
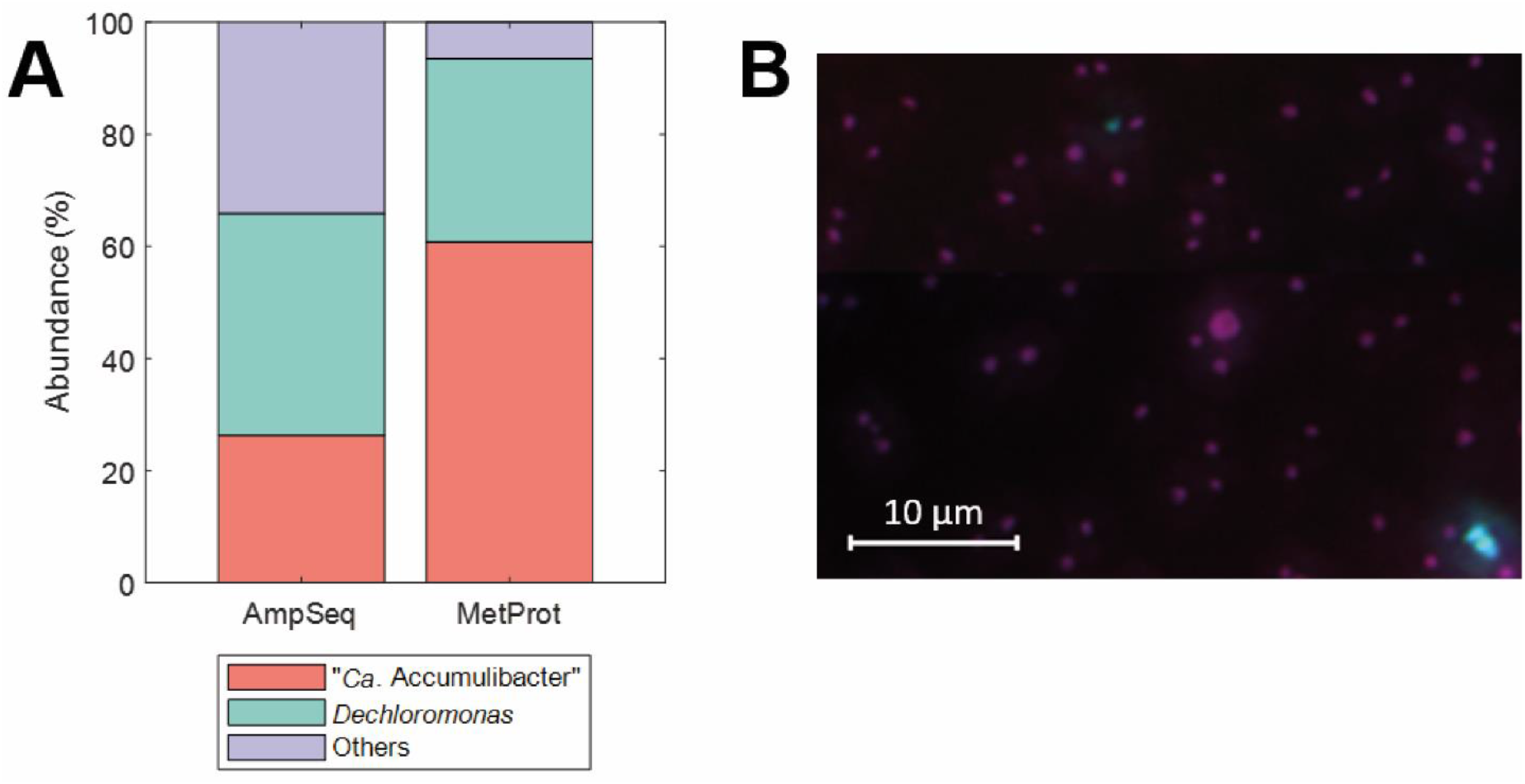
Microbial community analysis. *A:* Abundance of “*Ca*. Accumulibacter” and *Dechloromonas* based on 16S rRNA gene amplicon sequencing (AmpSeq) and metaproteomics (MetProt) based on identified peptides. *B:* Fluorescence *in-situ* hybridization (FISH) image of the PAO enrichment, with PAOmix probes (targeting “*Ca*. Accumulibacter”, in red), Dechmix probes (targeting *Dechloromonas*, in green) and EUBmix probes (targeting all bacteria, in blue). Magenta color represents the overlap of “*Ca*. Accumulibacter” (red) and eubacteria (blue); cyan color represents the overlap of *Dechloromonas* (green) and eubacteria (blue)

#### Analysis of ^13^C-labelled extracellular polymeric substances

After the enrichment achieved steady state conditions, the carbon source was switched from ^12^C-acetate to uniformly ^13^C-labelled acetate during approximately one SRT (8 days, 32 SBR cycles). Samples were taken at the end of each SBR cycle (*i.e*., at the end of the aerobic phase). EPS was extracted and analyzed. The protein and polysaccharide content of the extracted EPS was similar in all the samples and accounted for 55.9±3.7 and 6.2±0.7 % w/w of volatile solids of EPS. Note that these generally used measurements can have a significant bias (Felz et al. 2019). Mass spectrometric methods were used to determine the incorporation of ^13^C into polysaccharides and proteins. However, due to the harsh extraction conditions, other intracellular compounds may also be present in the extracted EPS as shown previously (Felz et al. 2016).

##### 3.1.1. ^13^C incorporation into sugars

In order to estimate the label incorporation in the extracellular polysaccharides, extracted EPS samples were hydrolyzed with HCl and the resulting sugars were analyzed by LC-HRMS. The labelling ratio of the sugars over the cultivation time is shown in Fig. 3. Label incorporation in sugars follow the same behavior as the theoretical dynamics due to excess sludge removal (solid line in Fig. 3). However, at the end of the experiment all analyzed sugars incorporated a higher amount of ^13^C than calculated based on the SRT (65 % enrichment). Moreover, two main trends can be distinguished. Glucose shows a clearly faster incorporation of the label when compared to the rest of the sugars. However, due to the extraction method, we cannot exclude presence of intracellular glycogen in our samples. In PAOs, intracellular glycogen is consumed and synthesized in every cycle, resulting in a high turnover rate, and therefore faster incorporation of the label (Mino et al. 1998). Thus, due to the possible presence of intracellular glycogen, glucose was excluded from the analysis extracellular sugars. The rest of the analyzed sugars present a common behavior, with lower dynamics, but still higher incorporation at the end of the experiment than expected on SRT dynamics only (solid line in Fig. 3).

**Fig. 3.**
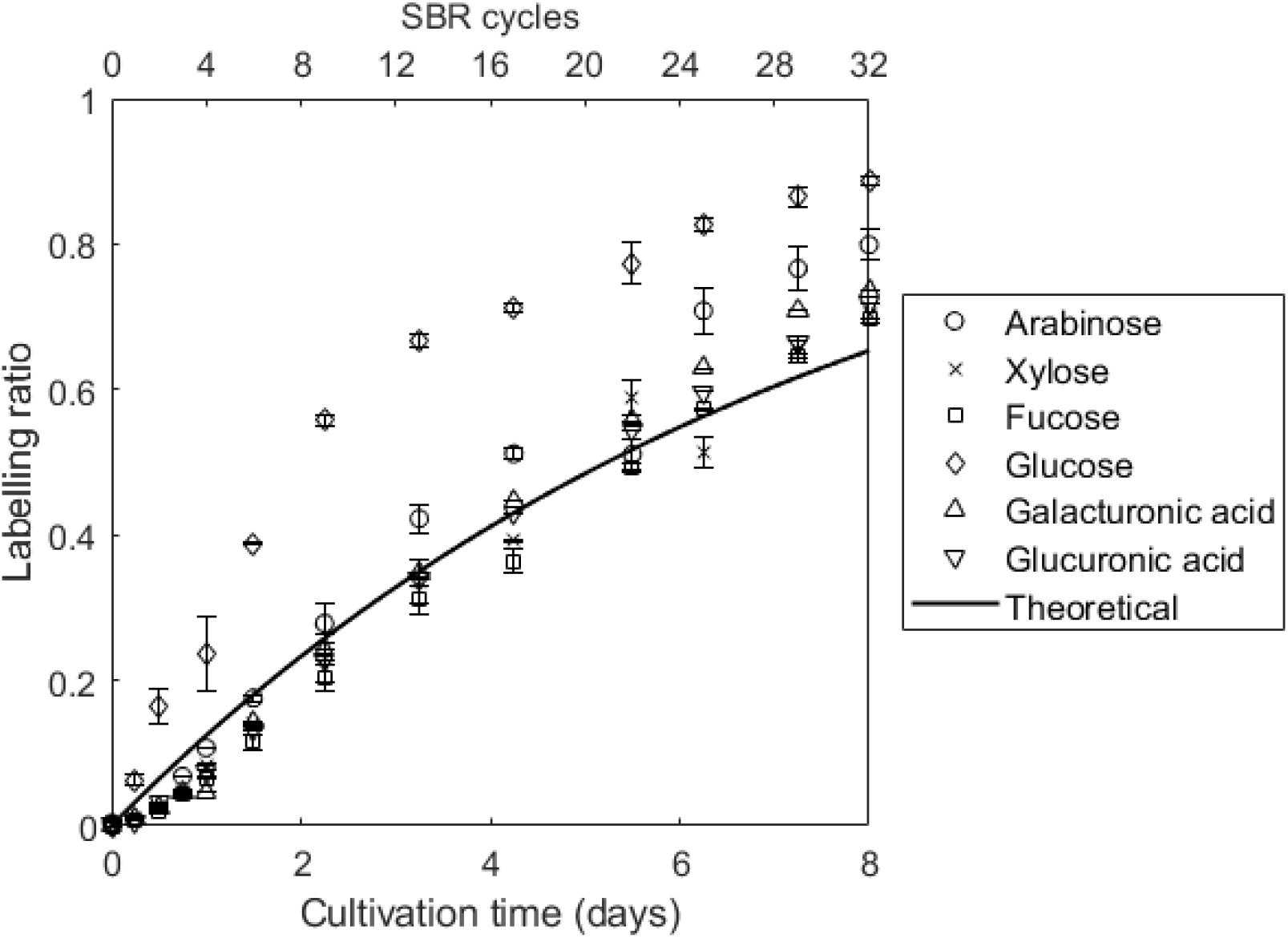
Labelling ratio for the several common monosaccharides over the different SBR cycles. Each point represents a sample at the end of a SBR cycle (top x-axis). The total duration of the experiment is shown in the bottom x-axis. The solid line represents the theoretical incorporation according to Eq. 1, with values of a = 1 and k = 0.132 d^-1^ (inverse of SRT). Error bars represent the standard deviation

A nonlinear regression analysis of the measured data was performed to estimate kinetic parameters as described by Eq. 1. This regression showed a high R^2^ value for all of the sugars (>0.97). Glucose showed a much faster turnover rate than the rest of the analyzed sugars. However, all the analyzed sugars exhibited a higher turnover rate than expected from the SRT (Table 1). This difference with the SRT indicates the presence of some degradation, compensated by a higher synthesis, and therefore, increased label incorporation.

**Table 1.** Turnover rate estimation of individual sugars. Calculated parameters of the regression analysis of time-dependent label incorporation in sugars according to Eq. 1. As a reference, the theoretical incorporation (solid line in Fig. 3) corresponds to values of a = 1 and k = 0.132 d^-1^ (inverse of SRT)

| Sugar | Asymptote, $a$ | Turnover rate, $k \text{ (days}^{-1}\text{)}$ | $R^2$ |
| --- | --- | --- | --- |
| Arabinose | 0.96 | 0.190 | 0.97 |
| Xylose | 0.96 | 0.151 | 0.98 |
| Fucose | 0.95 | 0.146 | 0.98 |
| Glucose | 0.98 | 0.326 | 0.99 |
| Galacturonic acid | 0.94 | 0.173 | 0.98 |
| Glucuronic acid | 0.96 | 0.158 | 0.99 |

##### 3.1.2. ^13^C incorporation into proteins

Extracted EPS samples were also analyzed by means of mass spectrometry based proteomics to monitor the ^13^C incorporation into the proteins. For a total of 782 proteins incorporation of ^13^C label could be detected. Detailed information about each individual protein can be further found in Table S1. The percentage of proteins showing labelling increased linearly over time. However, with increasing incorporation of ^13^C into the proteins, fewer proteins were identified. This is a known challenge in protein stable isotope probing due to broadening of peak isotope envelopes and the loss of fully unlabeled (native) peptide mass peaks in the spectra (Kleiner et al. 2021).

The labelling ratio of the most abundant stable isotope enriched cluster for all detected proteins was averaged, which value is shown in Fig. 4. Unlike the incorporation of label in sugars, label incorporation in proteins did not follow an exponential cumulative dynamic, but instead a sigmoidal incorporation. Label incorporation showed an initial slow incorporation up to approximately day 3 of the experiment. After this delay, the protein labelling ratio quickly increased until it started to approach an asymptotic value at the end of the experiment. The final average labelling ratio was higher than the value expected based on wash-out of biomass to maintain a constant SRT. Some proteins showed a longer or shorter delay in label incorporation, however, no specific trend was detected (Table S1**Table S1**). In order to compare to the behavior of proteins to sugars, protein data were also fitted using Eq. 1, and the derived parameters where in this case a = 0.92; k = 0.192 d^-1^; R^2^ = 0.88.

**Fig. 4.**
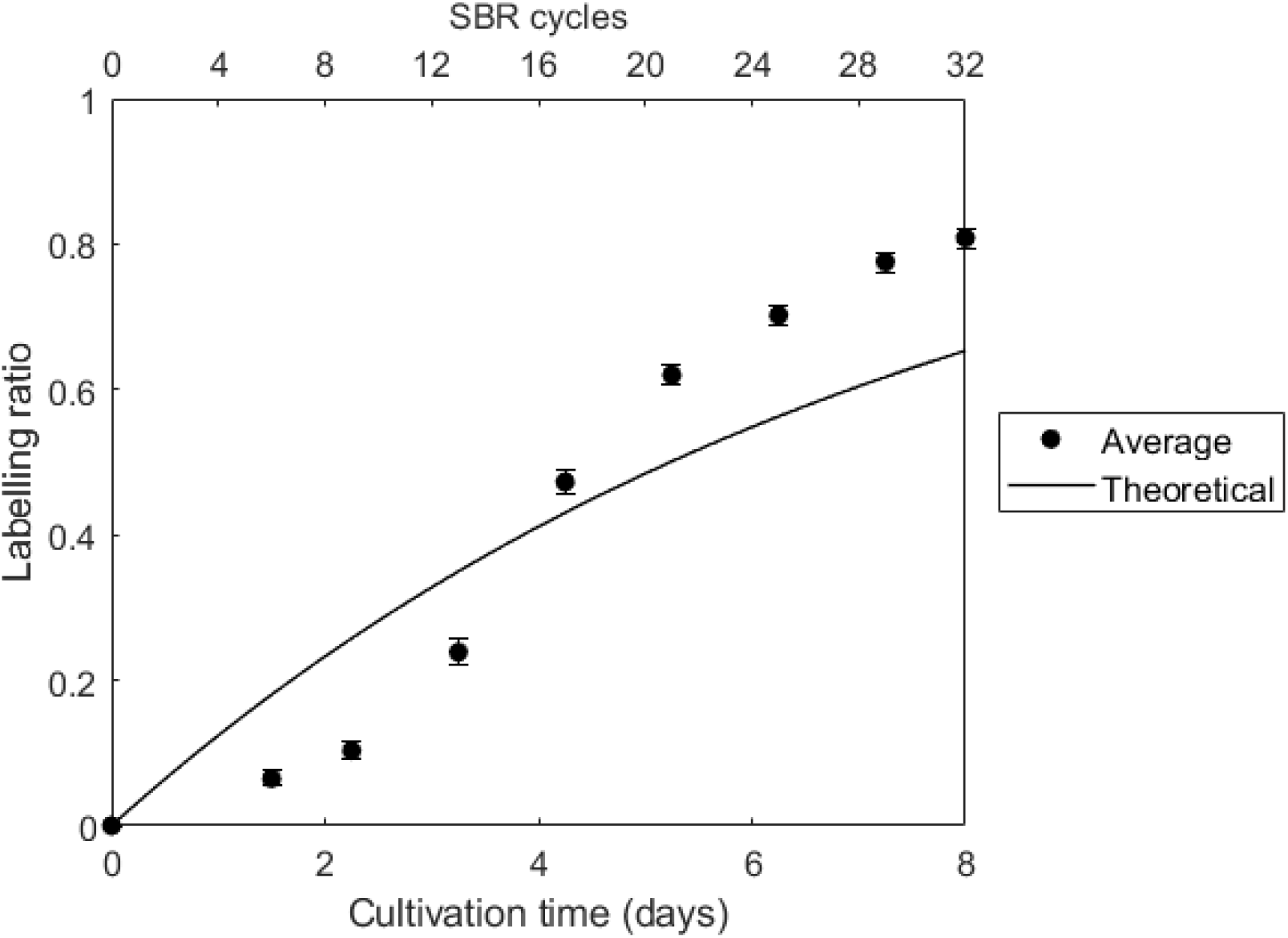
Labelling ratio for proteins over the different SBR cycles. Each point represents a sample at the end of a SBR cycle (top x-axis). The total duration of the experiment is shown in the bottom x-axis. The solid line represents the theoretical incorporation according to Eq. 1, with values of a = 1 and k = 0.132 d^-1^ (inverse of SRT). Error bars represent the 95 % confidence interval of the mean

Due to the harsh extraction method, intracellular proteins will also be present in the extracted EPS (Felz et al. 2016). In order to evaluate the differences of label incorporation between intracellular and extracellular proteins, a secretion-signal prediction tool (Almagro Armenteros et al. 2019) was used to distinguish intracellular from extracellular proteins. This tool recognizes the conserved amino acid sequences in the proteins that determine their secretion (Almagro Armenteros et al. 2019). The analysis revealed that 131 of the detected proteins (representing *ca*. 20 % of total protein signal intensity) have a signal peptide for secretion. This information was used to compare the label incorporation in secreted and non-secreted proteins (Fig. 5A). No difference was seen between both types of proteins as the average labelling ratio of both groups overlapped.

**Fig. 5.**
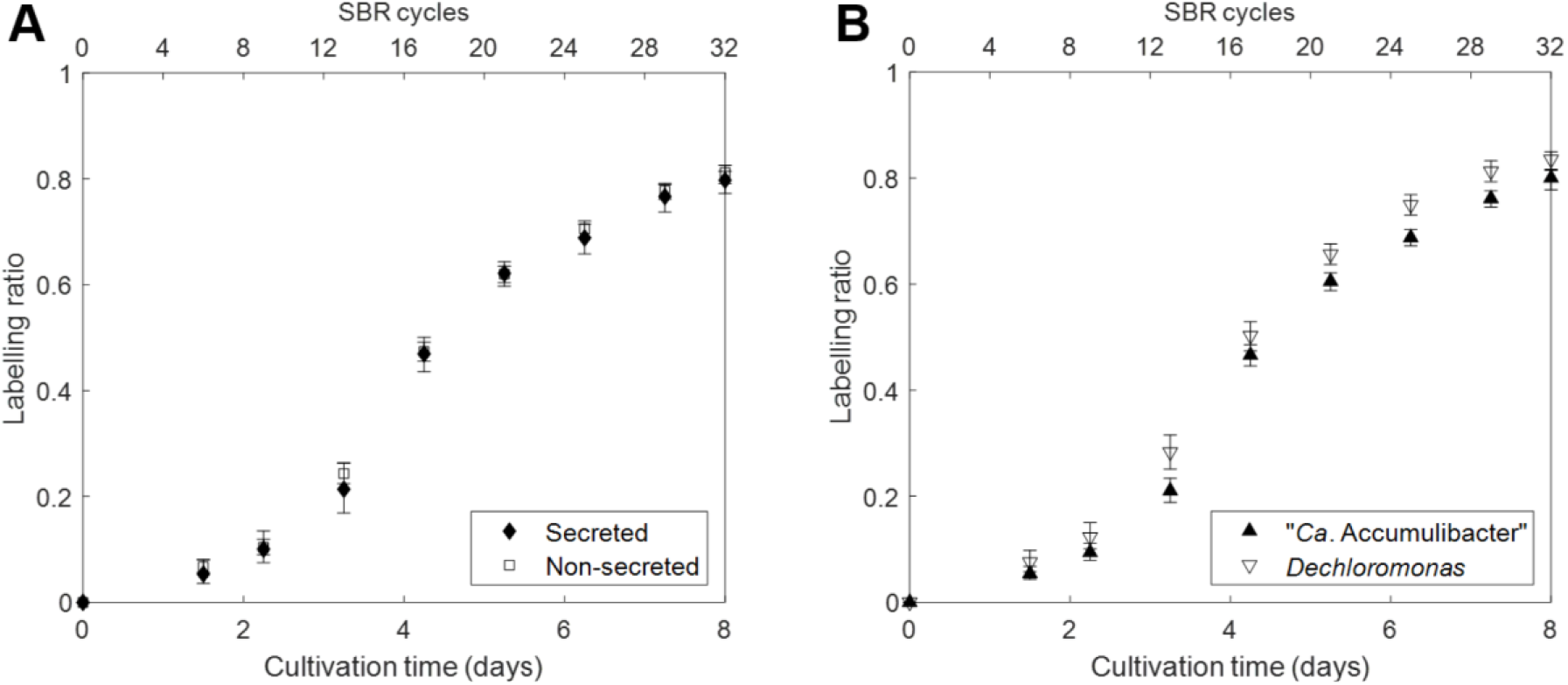
Labelling ratio observed for extracellular *vs*. intracellular (A) proteins or proteins from “*Ca*. Accumulibacter” *vs*. *Dechloromonas* (B) over the different SBR cycles. Each point represents a sample at the end of a SBR cycle (top x-axis). The total duration of the experiment is shown in the bottom x-axis. Error bars represent the 95 % confidence interval of the mean.

Additionally, proteins from the two PAO populations of *“Ca*. Accumulibacter” and *Dechloromonas* in the enrichment were analyzed separately to evaluate if they behaved differently (Fig. 5B). Proteins from both organisms showed the same trend with an initial delay. Although the labelling ratio in the proteins from *Dechloromonas* was slightly higher, no significant differences were observed.

## 4. Discussion

### 4.1. “*Ca*. Accumulibacter” and *Dechloromonas* showed a comparable incorporation behavior in the PAO enrichment

A SBR system simulating EBPR conditions was used to enrich for PAOs. The reactor conditions were similar to previous in-house work where the enrichments were dominated by ”*Ca*. Accumulibacter” (Welles et al. 2016; Rubio-Rincón et al. 2019). Periodic acetate and phosphate measurements and online conductivity and pH monitoring showed a stable and successful PAO enrichment. Anaerobic phosphate release coupled to acetate uptake was in the range of previously reported values (Oehmen et al. 2005; Welles et al. 2015), and the same as for a previous highly enriched culture of “*Ca*. Accumulibacter” (Guedes da Silva et al. 2020).

The microbial community analysis revealed that both “*Ca*. Accumulibacter” and *Dechloromonas* dominated the PAO enrichment. *Dechloromonas* has also been described as a PAO abundant in wastewater treatment plants (Petriglieri et al. 2021). The different community analysis methods (16S rRNA gene amplicon sequencing, proteome, FISH) gave different ratios for the two main species in the community. Analysis by 16S rRNA gene amplicon sequencing has been described to strongly underestimate the abundance of “*Ca*. Accumulibacter” (Albertsen et al. 2015; Kleikamp et al. 2022), but it can be useful to determine flanking populations. Therefore, we gave stronger credibility to FISH and proteomic results, which showed that these two PAOs formed the majority of the microbial community. *Dechloromonas* represented a substantial and stable fraction of the population based on protein counts, the exact reason for its selection in this enrichment culture is as yet unknown. *Dechloromonas* have been suggested to be denitrifying PAOs and play important roles in nitrogen cycle (Lv et al. 2014). However, no NO_3_^-^ was present in this cultivation and ATU was added to inhibit ammonium oxidation, which was only consumed for anabolic processes. Protein label incorporation pattern were highly comparable between *Dechloromonas* and “*Ca*. Accumulibacter”. Both PAOs have been described to share physiological characteristics (Kong et al. 2007), and both show a similar expressed proteome profile.

### 4.2. No difference in turnover was observed between extracellular and intracellular components

PAOs grow embedded in a matrix of EPS. EPS are a mixture of different biopolymers such as proteins or polysaccharides. In pure cultures and fast-growing organisms, EPS is mainly composed by polysaccharides (Sadovskaya et al. 2005). For slower growing organisms the EPS is more often composed of glycoproteins. Also for “*Ca*. Accumulibacter” (56 %), anammox bacteria (Boleij et al. 2018) and ammonium oxidizing bacteria (Yin et al. 2015) such glycoproteins have been described. EPS have been reported to be utilized by their producers or other neighboring microbes during nutrient limited conditions (Zhang and Bishop 2003; Pannard et al. 2016; Tomás-Martínez et al. 2022). In EBPR systems, carbon is rapidly taken up and stored intracellularly by PAOs, therefore no carbon source is available to other organisms present in the community and EPS could potentially be used as carbon source.

We examined the incorporation of ^13^C into proteins and polysaccharides, as constituents of EPS of PAOs. Due to the harsh extraction procedure, intracellular compounds can also be present in the extracted EPS (Felz et al. 2016). To differentiate intracellular and extracellular proteins, sequence analysis tools were used which revealed that *ca*. 20 % of the proteins corresponded to proteins containing a signal peptide for secretion. Although information about the contribution of intracellular components in the extracted EPS is limited, this value was higher than reported in the literature. For example, Zhang et al. (2015) detected 3 extracellular proteins out of 131 in activated sludge. The comparison of secreted and intracellular proteins showed no difference of label incorporation as both profiles overlapped (Fig. 5A). This indicates no difference between synthesis/degradation dynamics of proteins present in the extracellular or intracellular space, as ^13^C label incorporation follows the same dynamics. A higher degradation of a certain protein is balanced with a higher synthesis and therefore, faster label incorporation. Since the extracellular and intracellular proteins behaved the same, there was no indication for a significant higher turnover of extracellular proteins due to being used as carbon source by other bacteria present in the community.

Label incorporation into sugars did not reveal any unique labelling characteristics, with the exception of glucose, which was finally excluded from the analysis due to the potential intracellular glycogen present in the extracted EPS. The other sugars have been described previously as part of the EPS of aerobic granular sludge enriched with *“Ca*. Accumulibacter” (Weissbrodt et al. 2013; Felz et al. 2019). Overall, the turnover rate of polysaccharides (as combination of sugars) was lower than the one of proteins. Intracellular proteins showed a higher turnover rate than polysaccharides, indicating that extracellular components (such as polysaccharides) did not show a higher label incorporation (and therefore degradation) in the PAO enrichment. Recently, Arshad and colleagues demonstrated that there was no difference in the incorporation of label (^13^C or ^15^N) into the EPS and biomass of activated sludge (Arshad et al. 2021), which goes in line with the results obtained in our study.

Overall these result indicate that there is no faster turnover of extracellular polymers relative to intracellular biopolymers. In the literature it is often suggested that in biofilms a flanking community is growing on the extracellular polymers (Xu et al. 2022). However, this might be partly based on a too focused interpretation of previous studies. For instance, Zhang and Bishop (2003) only showed that extracted EPS can be degraded by starving bacteria. This is often used to state that EPS might have a preferential degradation by flanking populations. Using this *in situ* labelling approach, we identified equal turnover rates of intracellular and extracellular biopolymers. This indicates a general decay of biomass rather than a preferential consumption of EPS by the flanking populations.

### 4.3. SRT does not set the actual bacterial growth rate

Although extracellular and intracellular components did not show a difference in label incorporation, ^13^C was incorporated faster in proteins and sugars than what was expected based on the SRT. Based on the SRT (7.55 days), the average growth rate imposed to the experimental biosystem over the whole SBR cycle was 0.132 d^-1^ (μ_imposed_), as inverse of SRT. Based on the average turnover rates (Table 1, excluding glucose) of the different polymers, an average actual growth rate of 0.180 d^-1^ (μ_actual_) can be estimated, which would correspond to an SRT of 5.6 days. This difference between the imposed and the actual growth rate is the result of biomass decay. The actual growth rate is a combination of the imposed growth rate and the decay rate (k_decay_):

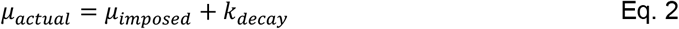

Based on the experimental observations a decay rate of 0.048 d^-1^ was calculated. Some models include decay rates for their growth simulations. For example, the general activated sludge model (Gujer et al. 1999), a decay rate for PAOs of 0.2 d^-1^ is included, which is an overestimation based on our observations.

This decay might be caused by different processes, such as, cell lysis, protein degradation or by the action of viruses or predators (*e.g*., other bacteria, protozoa). The observed decay corresponds to *ca*. 27 % of the actual growth rate of the system. This ratio of decay could explain the presence of *ca*. 7 % of flanking populations according to proteomic data (Fig. 2A). The label incorporation pattern in the different flanking populations could give more insights in this decay. However, due to the loss of resolution in the mass spectra, flanking populations could not be resolved in detail.

### 4.4. Incorporation of label in polysaccharides and proteins showed different dynamics

The incorporation of label in sugars and proteins showed a completely different behavior. For sugars the incorporation was according to the expected dynamics in line with most labelling studies (Cargile et al. 2004; Hong et al. 2012). On the other hand, incorporation in proteins appeared to show and initial delay comparable to a sigmoidal dynamics. A sigmoidal incorporation has previously been reported by (Taubert et al. 2011). However, in their case, the reason of this delay was a lag phase due to change of substrate. For environmental studies, however, often there are not enough data to determine whether the observed delay is also present in their experiments. On the other hand, in the study of (Martin et al. 2012) a short delay can be observed, but albeit at lower magnitude compared to our experiments. PAOs (and GAOs) present a unique metabolism involving the anaerobic-aerobic cycling of intracellular carbon storage polymers (*i.e*., glycogen and PHA). These polymers are initially non-labelled. In the first cycles the ^13^C would first build-up in these storage polymers and would then with a certain delay get incorporated in the microbial cell. It can however be calculated that this delay is only a few hours and not several days, Therefore the presence of these storage polymer pools cannot explain the observed delayed label incorporation in the proteins. Moreover if this was significant there would also be a delay in labelling of the sugars.

During the experiment, unlabelled yeast extract was present in low amounts in the medium. The main constituent of yeast extract are proteins, polypeptides and amino acids. The direct incorporation of these amino acids is likely not the cause of the observed sigmoidal incorporation curve for proteins. The yeast extract contributed only to *ca*. 1 % of the total COD of the feed, and its effect is marginal.

Another reason for this delay can be that synthesis of amino acids is more complex than sugar biosynthesis. It takes longer for bacteria to label their pool of amino acids. Moreover, the recycling of amino acids present in old proteins can also contribute to this delayed phenomenon. Bacteria can reutilize amino acids from misfolded or non-functional proteins through degradation by the proteasome or other less complex proteases (Elharar et al. 2014). The newly formed proteins would incorporate old non-labelled amino acids and newly synthesized labelled amino acids.

Finally, the employed approach to study ^13^C incorporation into proteins may not be sufficiently quantitative to resolve the low levels of incorporation in particular at early stages. Isotope patterns of metabolites (*e.g*., sugars) are easier to resolve, more homogenous, and therefore easier to measure and quantify. Nevertheless, all possible causes may contribute to the observed incorporation differences between proteins and sugars at the early stages, and deconvoluting their individual contributions will remain subject for further studies.

## Declarations

### Ethical approval

Not applicable

### Consent to participate

Not applicable

### Consent for publication

Not applicable

### Availability of data and material

The data generated and/or analyzed during the current study are included in this article and its supplementary material.

### Code availability

Not applicable

### Author Contributions

STM, EZ, MvL and DW planned the research based on intensive discussions among all the authors, especially with YL. STM and EW performed most of the laboratory work. MP conducted the proteomic analysis. FM performed the mass spectrometry analysis of sugars. STM and EZ interpreted the data with support of DW, MvL and YL. STM played major roles in drafting and writing the manuscript with input of DW, MvL and YL. All authors read and approved the manuscript.

## Acknowledgements

The authors would like to thank Samarpita Roy for providing the *Dechloromonas* sequences for the FISH analysis used in this research.

## Supplementary Material

**Fig. S1** Detailed microbial community composition. Relative genus-level microbial community distribution based on 16S rRNA gene amplicon sequencing (AmpSeq) and metaproteomics (MetProt) based on identified peptides. For AmpSeq, all OTUs contributing < 1 % are grouped as “Others”. For MetProt, all the groups cointributign < 0.5 % are represented by “Others”

**Table S1** Labelling ratio incorporation into the individual proteins over the different SBR cycles. Functional and taxonomic annotation and subcellular location prediction is shown for each protein

