## Supplementary figures and images for "Turnover of the extracellular polymeric matrix in an EBPR microbial community"

### Fig. S1

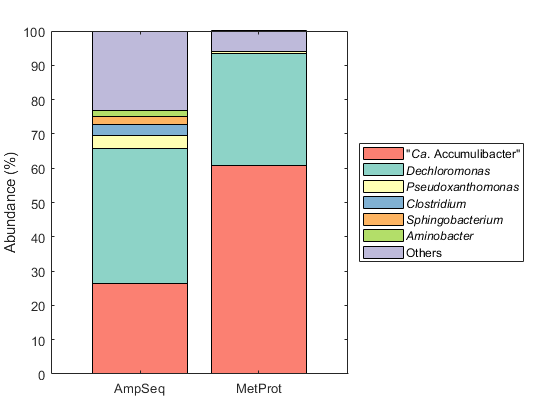
